# A *De Novo* Genome Assembly, Gene Annotation, And Expression Atlas For The Monarch Butterfly *Danaus plexippus*

**DOI:** 10.1101/2020.09.19.304162

**Authors:** José M. Ranz, Pablo M. González, Bryan D. Clifton, Nestor O. Nazario, Pablo L. Hernández-Cervantes, María J. Palma-Martínez, Dulce I. Valdivia, Andrés Jiménez-Kaufman, Megan M. Lu, Therese A. Markow, Cei Abreu-Goodger

## Abstract

The monarch butterfly epitomizes insect biodiversity decline. Understanding the genetic basis of the adaptation of the monarch to a changing environment requires genomic and transcriptomic resources that better reflect its genetic diversity while being informative about gene functionality during life cycle. We report a reference-quality genome assembly from an individual resident at a nonmigratory colony in Mexico, and a new gene annotation and expression atlas for 14,865 genes, including 492 unreported long noncoding RNA (lncRNA) genes, based on RNA-seq data from 14 larval and pupal stages, plus adult morphological sections. Two thirds of the genes show significant expression changes associated with a life stage or section, with lncRNAs being more finely regulated during adulthood than protein-coding genes, and male-biased expression being four times more common than female-biased. The two portions of the heterochromosome *Z* display distinct patterns of differential expression between the sexes, reflecting that dosage compensation is either absent or incomplete –depending on the sample– in the ancestral but not in the novel portion of the *Z*. This study represents a major advance in the genomic and transcriptome resources available for *D. plexippus* while providing the first systematic analysis of its transcriptional program across most of its life cycle.

## INTRODUCTION

The monarch butterfly *D. plexippus* is known for its spectacular annual migrations across North America (1,2). Nevertheless, the well documented population decline in this species is of increasing concern. For example, the census of monarchs across overwintering sites in Mexico during the period 2013-2014 was just ∼10% of the last 20-year average (3). This trend is explained by multiple factors, including a loss of the Mexican overwintering habitat due to illegal logging (3), agricultural practices that reduce the populations of the most suitable milkweed *Asclepias* species for female oviposition (Supplemental Fig. S1)(4), and their replacement by non-optimal alternative hosts such as milkweed invasive species *Gomphocarpus physocarpus* (5).

Central to the future of the natural populations of *D. plexippus* is an accurate understanding of its unique host specificity and potential for host shifts, the basis for its pesticide and parasite resistance, and other uncommon aspects of its biology in comparison to other Lepidoptera and insects in general. An increasing number of studies have aimed to address some of these questions (6-8), some of them developed in a more comparative genomic context (9,10). Unfortunately, however, the monarch research community still relies upon a single *D. plexippus* genome assembly, which was only recently upgraded to reference-quality standards (7,11). Equally important, and despite relevant efforts to gain insights into gene functionality in *D. plexippus* (8,12-15), a comprehensive expression atlas during the life cycle of this species is still missing. Thus, there are fundamental gaps in knowledge for different sets of genes such as for long noncoding RNA genes (lncRNAs). These are virtually absent from the gene annotation *D. plexippus* (7), despite being increasingly recognized due to their functional and phenotypic importance (16-18).

Given the urgency associated with the rapid decline of *D. plexippus* populations, development of enhanced genomic and transcriptomic resources more reflective of the underlying genetic variation of the species and more informative about basic aspects of its biology and development is crucial. Here, we have generated a *de novo* genome assembly, a *de novo* gene annotation, and a transcriptome atlas that includes lncRNA genes, over 14 different biological conditions representing different life stages or body parts from adult males and females from a non-migrant colony of Mexico. With these resources, we have reexamined the genome organization of *D. plexippus* in relation to other Lepidoptera, and provided a rich portrait of transcriptome patterns during the life cycle of this species that address diverse aspects such as sex-biased gene expression and dosage compensation.

## MATERIAL AND METHODS

### Butterfly husbandry

Newly hatched larvae of *D. plexippus* were collected from wild *Asclepias curassavica* on the campus of the National Laboratory for Genomics of Biodiversity in Irapuato, Guanajuato, México. The early first instar caterpillars were placed in individual vials and fed with fresh *A. curassavica* leaves *ad libitum* on a 12-12h light-dark cycle at 25°C and ∼50% relative humidity until adults emerged. All stages were precisely identified measuring head capsules left after molting.

### Genomic DNA extraction and sequencing

A two-day-old pupa was fast frozen in liquid nitrogen and preserved at -70°C until DNA extraction. Genomic DNA was extracted with the Blood and Cell Culture DNA Kit (QIAGEN). All equipment was cleaned with DNAaway (Thermo-Fisher Scientific) prior to grinding the specimen in a mortar kept cold with liquid nitrogen. Powdered pupal tissue was incubated with RNAse-A and Protease for 2h while gently rocking. DNA was purified by gravity flow with QIAGEN Genomic-tips, precipitated with isopropanol, and washed twice with cold 70% ethanol. All centrifugation steps were performed in 15 ml tubes in a pre-chilled centrifuge at 4°C. The DNA pellet was resuspended in QIAGEN EB buffer overnight. One μg of unsheared genomic DNA was saved for Illumina PE-150 sequencing in a HiSeq 4000 instrument over one lane. The remaining genomic DNA was sheared with blunt-end needles as reported (19); except for 20 pumps with a 21 gauge 1.5’’ blunt end needle, followed by 10 pumps with a 24 gauge 1.5’’ blunt end needle (Jensen Global, Santa Barbara, CA). Ten μg sheared DNA was used to make the SMRT-bell template library following manufacturer specifications. The library was size selected (15-80 kb) using the Blue Pippin size selection instrument (Sage Science) and then sequenced on six SMRT cells (one cell at 2 pM and five cells at 6 pM) using 1M v2 chemistry on a PacBio Sequencing Sequel instrument with a 10 hr movie time. Concentration and purity of all the genomic DNA submitted for sequencing was determined using a Qubit v2 fluorometer (Life Technologies) and an 8000 NanoDrop (Thermo Scientific), respectively. All genome sequencing was performed at the UCI Genomics High-Throughput Facility.

### Total RNA extraction and sequencing

Fourteen specific stages and anatomical parts were RNA-sequenced: 1^st^ (pools of six), 3^rd^ and 5^th^ instar larvae; day 1, 3, 5, 7 and 9 pupae; 2-day-old adult female and male thoraces, heads, and abdomens. Adult individuals were anesthetized in a cage at -20C for five minutes, sexed, and the wings removed, after which they were dissected into the indicated anatomical sections. All samples were fast-frozen in liquid nitrogen and preserved at -70°C until RNA extraction. With the exception of 1^st^ instar larvae and adult heads, which were mechanically homogenized in TRIzol using Teflon homogenizers, the rest of sample types were weighed after pestle homogenization in a ceramic mortar using liquid nitrogen, adjusting for sample quantity prior to be stored in TRIzol. Total RNA was subsequently extracted using Direct-zol RNA MiniPrep extraction kit (Zymo Research) according to manufacturer recommendations, except for all pupa samples and adult abdomens which were first extracted with TRIzol also following manufacturer recommendations and then purified with Direct-zol columns. RNA yield, purity, and integrity were evaluated with conventional methods, using a Qubit 2.0 Fluorometer, a NanoDrop 8000 Spectrophotometer, 1% agarose gels, and RNA 6000 Pico and RNA 6000 Nano kits –depending on the experiment– with a BioAnalyzer 2100 (Agilent Technologies Inc.). Libraries for each sample type were prepared using the TruSeq Stranded Total RNA Library Prep Kit (Illumina), multiplexed and 75 bp paired-end sequenced over 4 lanes on an Illumina NextSeq 500 Sequencing System at the sequencing core facility at LANGEBIO. Aliquots from samples of pooled individuals belonging to the same broadly defined developmental stage (*Lpool* = 1^st^, 3^rd^, and 5^th^ instar larvae; Ppool = 1, 3, 5, 7 and 9-day pupae; *Mpool* = heads, abdomens, and thoraces from adult males; *Fpool* = heads, abdomens, and thoraces from adult females) were mixed equimolarly. For these pooled samples, non-poly(A) enriched stranded libraries were constructed using the TruSeq Stranded Total RNA Library Prep Kit (Illumina) and the Ribo-Zero Gold Set A kit (Epicenter). These libraries were subsequently multiplexed and 100 bp paired-end sequenced over one lane on an Illumina HiSeq 2500 instrument at the University of California Irvine Genomics High Throughput Facility.

### *De novo* genome assembly construction

We generated different assemblies to be used in different analyses or to be associated with different stages part of the same computational pipeline. Illumina paired-end reads were trimmed and filtered out for low-quality base calls and undesired adapter presence using Trimmomatic v.0.35 (20) and FastQC 0.11.5 (21), and used to generate an assembly with Platanus v.1.2.1 (22), which can accommodate any residual heterozygosity, using default parameters. The quality of the Platanus assembly was confirmed upon finding that the mapping back rate, *i*.*e*. the percentage of reads aligned against the constituent collection of contigs, was ∼99.4%. A *k*-mer analysis was performed to estimate the level of heterozygosity in our sequenced sample and to recalculate the genome size, as a control, of *D. plexippus*, using GenomeScope (23).

A draft genome assembly based on PacBio raw sequencing reads was used using Canu v1.6 (24) specifying a genome size of 250 Mb, a corrected error rate of 0.045, a raw error rate of 0.3, a minimum overlap length of 500 nt, and a minimum read length of 1000 nt. The resulting assembly featured an NG50 = 3.3 Mb (NG50 refers to the length of the smallest contig added to cover 50% of all nt estimated in the genome; (25)), a total size of 458.6 Mb, and an error rate upon self-correction of 0.045. This intermediate assembly was subsequently polished through four rounds of Pilon v1.22 (26) using the alignment information from Illumina trimmed reads generated with bwa v0.7.17-r1188 (27). Redundancy minimization was performed with HalploMerger2_20180603 (28) using default parameters except for splitting the target in fast files of 5×10^6^ nt instead of 5×10^7^ nt and with a query size of 160×10^6^ nt instead of 160×10^7^ nt. Then, FinisherSC (29), along with MUMmer v4.0.0beta1 (30), was used to upgrade the haploid assembly using all raw PacBio reads (NG50 = 5 Mb, total size = 434.9 Mb). At this stage, we polished the new expanded diploid assembly again through five additional rounds of Pilon v1.22, followed by HalploMerger2_20180603, to generate the final haploid collection of contigs. These contigs were scaffolded with RaGOO (31) using one of the other assemblies generated as a reference (Quickmerge in Supplemental Table S1). This alternative assembly was obtained in the course of our attempts to enhance contiguity and resulted from merging our polished DBG2OLC assembly, which combined the Illumina-based Platanus assembly and raw PacBio reads, with our polished Haplomerger2 canu-derived assembly.

Quality metrics of the selected and non-selected assemblies, as well as key intermediate assemblies generated through different approaches, were extracted using the script NX.pl (https://github.com/YourePrettyGood/RandomScripts/blob/master/NX.pl). Genome assembly completeness was assessed by calculating different mapping back rates of sequencing reads from 72 Illumina genomic DNA sequencing libraries (6) that were considered suitable (see below). Read mapping was done with bwa v0.7.17-r1188 under the “-h 99999” setting, and the different rates calculated using the counts for mapped, properly paired, and total reads as obtained with SAMtools v1.9 (32). Gene-level completeness was evaluated through CEGMA v1.0 (33), and BUSCO v2.0.1 and BUSCO v4.0.5 (34), using the gene sets of Endopterygota_odb9 (n = 2,442) and Lepidoptera_odb10 (n = 5,286), respectively. Lastly, differences in scaffolding between DpMex_v1 and Dpv3 were determined with RaGOO (31) using the former as a reference, which allowed the identification of chimeric contigs in Dpv3. Briefly, if a Dpv3 scaffold aligns against two different DpMex_v1 contigs over at least 10 kb in each case, and the span covered of these contigs was in both cases greater than 100kb and 5% of the contig span, the Dpv3 scaffold was dubbed as chimeric.

### Repeat annotation

*Ab initio* repeat modelling was done with RepeatModeler v.1.0.11 (35). The filtered RepeatModeler database was combined with consensus Lepidopteran repeats found at Dfam_Consensus-20170127 (36) or RepBase-20170127 (37) databases. The global set of repeats was used to feed RepeatMasker v.4.0.7 (38) to softmask the final polished genome assembly.

### Gene annotation

Funannotate v1.5.3 docker image (39) was used to train Augustus v3.2.3 (40), predict gene models, and perform functional annotation. As input for optimizing the performance of Augustus v3.2.3 (40), funannotate used 2,404 PASA v2.3.3 gene models (41). To obtain this training gene model set, transcripts were de novo assembled with Trinity v2018-2.8.3 under settings --SS_lib_type RF (42), using all poly(A) RNA-seq paired reads after adapter removal with Trimmomatic v0.32 (20). These transcripts were aligned to the genome under PASA using BLAT v36 (43), obtaining a first set of gene models. The 500 longest non-redundant ORFs associated with the PASA gene models were used to train TransDecoder v5.2.0 (44). Then the gene models were selected according to their abundance as estimated by Kallisto v0.44.0 (45) under settings --rf-stranded using the Trinity normalized reads. Ultimately, BRAKER v2.0.3b (46) trained Augustus with the retained gene models.

For gene prediction, funannotate aligned mRNAs and proteins from the previous annotation (official gene set 2, OGS2)(7) with minimap v2.14-r883 (47) under settings -ax splice --cs -u b -G 3000, and Diamond blastx v0.8.22 (48), respectively. Protein alignments were further refined by funannotate, including 3 kb upstream and downstream of the region of alignment, and subsequently executing Exonerate v2.4.0 (49). Additionally, funannotate parsed the introns supported by alignments of poly(A) RNA-seq reads generated with HISAT v2.1 (50) under settings --rna-strandness RF --max-intronlen 10,000. This combination of hints (protein alignments, transcript alignments, and intron locations) was used by Augustus to predict a second set of 16,756 gene models. Of them, 9,695 were dubbed as highly supported, *i*.*e*. had more than 90% of their model supported either by intron hints, transcript alignments, or protein alignments. GeneMark-ET v4.35 (51), under settings --max_intron 3,000 --soft_mask 2,000, was also run independently to predict a third set of gene models but only relying on intron hints.

The PASA, Augustus highly supported, Augustus not highly supported, and GeneMark prediction sets were combined by EVidenceModeler (52), assigning them 10, 5, 1, and 1 relative weights, respectively. The predictions were further filtered by removing genes shorter than 50 aa in length, or that had high sequence similarity (diamond blastp --sensitive --evalue 1e-10) to the repeat database included in funannotate, or that had more than 90% of the model intersecting regions masked by RepeatMasker. The filtered set of gene models was updated in order to include UTR information by two executions of the PASA annotation comparison using the Trinity transcripts and filtering gene models according to transcripts per million as calculated by Kallisto. Alternative transcripts were only kept if they were at least 10% as highly expressed as the most highly expressed transcript per gene.

Non-coding genes were annotated with the following tools: tRNA genes, tRNAscan-SE v.2.0 (53); rRNA genes, RNAmmer v.1.2 (54); and for a variety of other RNA genes, Infernal v1.1.1 (55). Specifically, for miRNA-encoding genes, we used BLASTn to locate the most recent annotation of these genes (56). Lastly, FEELnc (57) classified lncRNAs from the transcripts assembled by StringTie v1.3.2d (58), and considering protein-coding predictions described above.

### Homology identification

The preliminary new set of protein-coding genes as from funannotate was then used by OrthoFinder v2.2.6 (59) under the settings –S diamond –M msa to establish orthologous calls across protein sets from 7 other species, which were retrieved either from NCBI or lepBase (Supplemental Table S7). Only the longest predicted protein per gene model was used in the analysis. Orthogroups with other lepidopterans and *D. melanogaster* were identified independently for our gene predictions and the annotation of the previous assembly, *i*.*e*. OGS2. Also, when identifying gene correspondence between our predictions and OGS2, all other species were excluded from the input to OrthoFinder. 1-to-1 orthologs were grouped in microsynteny blocks by DAGchainer (60) under default parameters. The software Circos (61) was used to graphically represent the chromosomal mapping of microsynteny blocks between lepidopterans.

### Sex-dependent sequence coverage analysis

Illumina genomic DNA sequencing data from 80 *D. plexippus* individuals were retrieved (6). Supplemental Table S9 lists their GenBank SRA accession numbers. Sequencing reads from each sample were aligned against the DpMex_v1 assembly using bwa v0.7.17-r1188 under the “-M” option. Contig median coverage was calculated using mosdepth (62). We calculated a normalized contig coverage for each sample as the contig scaffold coverage divided by a weighted average, according to the total number of reads mapped to each contig, of the median contig coverage. Five samples (SRR1548577, SRR1549538, SRR1552003, SRR1552104, and SRR1552110) were filtered out due to having less than an average coverage of 5, which was coincidental with an unusual distribution of their normalized coverage relative to the rest of samples. To estimate the male:female (M:F) coverage ratio, we averaged the normalized coverage per contig per sex as reported (9). Further, the cumulative fraction of the Z chromosome covered for at least a given normalized coverage value was plotted to explore the presence of outlier samples. Two samples labelled as female (SRR1552102 and SRR1552103) stood out as they had a normalized coverage above 0.98 for more than 50% of the heterochromosome *Z*, which is similar to the typical distribution for male samples. Similarly, one sample labelled as male (SRR1552006) had a normalized coverage distribution that resembled that of females (Supplemental Fig. S12). Lastly, two additional samples (SRR1548506 and SRR1549526) exhibited highly heterogeneous median coverage among contigs. These five samples were also excluded from further analyses.

### Genomic clustering of developmental genes

For *D. plexippus*, we used our newly generated assembly (scaffold N50 ∼8 Mb; DpMex_v1) and gene annotation (OGS1_DpMex). For other species, we used the assemblies and annotations available in Ensembl (http://ensemblgenomes.org/): *B. mori*, ASM15162v1 (scaffold N50 ∼8 Mb); and *D. melanogaster*, BDGP6 (scaffold N50 ∼25 Mb). To compare the relative location of genes that belong to particular clusters between *D. plexippus* and *D. melanogaster*, we used information from 1-to-1 orthologs obtained with OrthoFinder as well as the outcome of TBLASTN reciprocal best hit searches between *D. melanogaster* and *D. plexippus* (63). Homology searches were done: for *D. plexippus*, by running in-home shell scripts in the UCI High Performance Computer cluster; for *B. mori* and *D. melanogaster*, using the BLAST tools available in Ensembl. Clustal omega was used to visually inspect protein sequence alignments and resolve cases of potential gene model fragmentation (64). Regardless the presence of intervening genes, we considered the existence of clustering if the distance between two relevant genes was ≤0.1% of the total assembly size, *i*.*e. ∼*250 kb in *D. plexippus* and 120 kb in *D. melanogaster* (for the latter, only considering the euchromatic fraction). When appropriate, *e*.*g*. to confirm gene duplications and deletions, we extracted genome sequences from genome assemblies and performed local alignments using PipMaker (65).

### RNA-Seq quality control and alignment

The raw sequencing data in fastq files were preprocessed with fastqc (21) and multiqc (66) to verify that the sequencing was of sufficient quality; all files passed visual inspection. Reads were aligned to the genome based on a two-pass strategy using STAR v2.7.3a (67). The genome was first indexed for STAR including the exons from the GFF annotation file. During the first pass of alignments, only the Splice-Junction files were stored. The complete set of Splice-Junction files were used during the second pass to inform the final alignments. Non-default parameters used during both passes are: --outFilterMultimapNmax 500, --outFilterMismatchNoverReadLmax 0.1, --alignIntronMin 5, and --alignIntronMax 20000. Adapter sequences were trimmed during alignment, using the parameter -- clip3pAdapterSeq AGATCGGAAGAGCACACGT AGATCGGAAGAGCGTCGTG.

### Gene-level expression quantification

The annotation file was first processed to remove overlaps, using the R package GenomicRanges (68). As protein-coding and lncRNAs were our focal interest (first set), they were considered separately from rRNA, tRNA and miRNAs (second set). Any genomic coordinate overlaps between the second and the first gene types were deleted from the first. All the remaining coordinates in both sets were collapsed at the exon level. Introns were determined as the spaces left between collapsed exons for every gene. The resulting annotation was used as the input for featureCounts (69) in order to determine separate exon and intron gene expression counts for each library. Non-default parameters were: largestOverlap=TRUE, fraction=TRUE, strandSpecific=2, and isPairedEnd=TRUE. Most libraries presented a high percent of rRNA counts, but low counts assigned to introns and tRNAs, which were all discarded before further processing (Supplemental Table S11). Exonic gene-level expression for proteincoding, lncRNA, and miRNA genes were stored as raw counts (Supplemental Table S12).

### Differential expression analysis

Raw counts were further processed using the edgeR package (70). Each type of sample (*e*.*g*. L1) was assigned to a distinct factor level. Only genes with ≥1 count-per-million (CPM) in at least two samples were kept. Normalization factors were calculated with the calcNormFactors function and the TMM method (71). Normalized log_2_CPM, as well as fragments-per-kilobase-per-million (FPKM) expression, values were saved for subsequent analyses. Multi-dimensional Scaling plots were used to determine the relationship between samples and grouping of replicates. During analyses, several samples from individuals were determined to have a strong male-specific expression profile. These samples (L5.1, P1.1, P3.1, P5.1, P7.1, P7.2, P9.1, MT.1, MT.2) were assigned to an extra “male” batch factor. Negative binomial dispersion values were calculated, used to fit generalized linear models, and to test for differential expression with glmTreat (72). This approach tests for differences in expression that are significantly higher than a threshold, in this case a fold-change of 2. Finally, to select differentially expressed genes, a 0.05 False-Discovery Rate (FDR) was used using the Benjamini-Hochberg multitest correction method (73). In some analyses, a less strict likelihood ratio test was also performed to find fold-changes significantly higher than 0. For both approaches, 24 differential expression contrasts were chosen to represent individual samples and transitions across the atlas (Supplemental Table S13).

### Transcriptional network and clustering

Weighted Gene Correlation Network Analysis (WGCNA) (74) was used to generate a transcriptional network, considering all 14,865 genes that were also used for differential expression. The library size normalized log_2_CPM expression values were used. The pooled samples sequenced at the UCI facility (Supplemental Table S11) were first removed to avoid correlations between different sequencing facilities and library prep methods affecting the network. As recommended by the package authors, a range of soft-threshold values were explored, and 14 was selected to optimize the fit of the network to a scale-free topology. A topological overlap similarity matrix was calculated, preserving the sign of the correlations. Hierarchical clustering with the average agglomeration method, and a dynamic tree cutting procedure were used to obtain gene clusters. To allow for relatively smaller clusters, the minimum module size was set to 10, which resulted in 25 clusters.

### GO enrichment analysis

For each group of genes resulting from differential expression or network clustering, a test for enrichment of GO terms was performed using clusterProfiler (75). All GO terms assigned in our annotation were considered, as well as all their ancestor terms. During each enrichment test, GO terms with less than 5 or more than 500 genes were ignored. Although GO terms are not independent due to their hierarchical nature, multiple-testing correction using the q-value method was performed (76). A q-value cutoff of 0.2 was used as a threshold for GO term enrichment.

### Statistical analyses and figure plotting

Most statistical analyses were executed primarily in R (77). Figure plotting was also largely performed using base R functions, as well as those included in the R packages beanplot (78), ggplot2 (79), ggVennDiagram (80), and pheatmap (81). The packages cowplot (82) and gridGraphics (83) were additionally used for the layout of some multi-panel figures.

## RESULTS AND DISCUSSION

### *De novo* genome assembly

A single pupa of *D. plexippus* was collected at Irapuato, Mexico, and sequenced using both Illumina PE-150 and PacBio Single Molecule Real-Time (SMRT) technologies under strict conditions to prevent contamination from unintended species (Material and Methods; Supplemental Text S1 and Fig. S2). A total of ∼97.3 Gb of raw Illumina data were generated and filtered resulting into ∼72.4 Gb of high quality trimmed reads, representing a 255x sequence coverage –assuming a genome size of 284 Mb (84). In parallel, and through PacBio, we obtained a sequencing output with a 193x sequence coverage (subreads > 1 kb only) and an NR50 of 22.6 kb (the median read length above which half of the total coverage is contained; Fig. S3), a value higher than that associated with recently published reference-quality genome assemblies of *D. melanogaster* (85). To generate a *de novo*, reference-quality genome assembly, we adopted different computational strategies that ultimately led to a limited set of genome assemblies, from which one was selected (Supplemental Table S1; Fig. S4). This assembly exhibits enhanced contiguity (Fig. S5A), encompassing 108 contigs polished at different stages with the Illumina sequencing data, with an additional contig (Sc0000031) very likely representing a different haplotype for Sc0000030 (Supplemental Text S2 and Fig. S6). In total, 78 contigs were merged into 36 scaffolds (see below). The final assembly, DpMex_v1, features a scaffold N50 of 8.16 Mb (Table 1), and a heterozygosity of 2.15% (Fig. S5B). Further, we evaluated the genome assembly and gene set prediction completeness. In the first case, we mapped DNA Illumina sequencing reads from 72 samples back onto the DpMex_v1. The global fraction of mapped sequencing reads was 93.6% and that for those discordantly mapped of 3.67%, both very similar to those obtained against the Dpv3 assembly (6) (Supplemental Table S2). Next, gene-level completeness was ascertained using the nearly-universal set of single-copy genes using BUSCO v4.0.5 (34). We recovered 98% complete BUSCOs in the Lepidoptera gene set (lepidotpera_odb10, n=5826), with an additional 0.5% in multiple copies (Table 1). Together, these results provide a new monarch genome assembly, DpMex_v1, which is highly contiguous and virtually complete.

**Table 1.** Salient features of the *D. plexippus* genome assembly obtained here compared to other relevant ones.

|  | *D. plexippus* | *D. plexippus* | *D. plexippus* * | *M. cinxia* * | *H. melpomene* * | *B. mori* * |
| --- | --- | --- | --- | --- | --- | --- |
| Assembly Identifier | DpMex_v1 | Dpv4 | Dpv3 | v1.0 | Hmel2.5 | ASM15162v1 |
| Span (Mb) | 248.571 | 248.676 | 248.564 | 389.9 | 275.246 | 480.5 |
| GC content (%) | 32.2 | 31.6 | 31.6 | 32.6 | 32.8 | 37.7 |
| Contigs |  |  |  |  |  |  |
| Number | 108 | 10,682 | 15,441 | 48,180 | 3,126 | 88,673 |
| N50 (kb) / NumN50 | 3,940 / 21 | 111.0 / 548 | 63.6 / 906 | 14.1 / 7366 | 328.9 / 214 | 1.6 / 8,076 |
| Scaffolds |  |  |  |  |  |  |
| Number | 65 | 4,115 | 5,397 | 8,261 | 332 | 43,463 |
| N50 (kb) / NumN50 | 8,158.1 / 13 | 9209.9 / 12 | 716.0 / 101 | 119.33 / 970 | 14,309.0 / 9 | 4,008.4 / 38 |
| N90 (kb) / NumN90 | 3,498.3 / 30 | 5644.642 / 25 | 160.5 / 366 | 29.60 / 3396 | 9,045 / 19 | 61.1 / 258 |
| Longest (Mb) | 16.34 | 15.62 | 6.2 | 0.67 | 18.1 | 16.2 |
| Ns (%) | 0.002 | 2.73 | 2.7 | 7.42 | 0.4 | 10.4 |
| CEGMA (n=248) † | C: 93.15%<br>P: 96.8% | C: 92.34%<br>P: 96.8% | C: 90.3%<br>P: 96.8% | C: 68.55%<br>F: 86.7% | C: 88.71<br>P: 96.77% | C: 76.6%<br>P: 96.8% |
| BUSCO (n = 5286) † | C: 98.5%<br>D: 0.5%<br>F: 0.5% | C: 98.3%<br>D: 1.4%<br>F: 0.5% | C: 98.5%<br>D: 1.6%<br>F: 0.5% | C: 91.8%<br>D: 0.5%<br>F: 3.7% | C: 98.8%<br>D: 0.6%<br>F: 0.3% | C: 95.3%<br>D: 0.3%<br>F: 1.7% |

\* Retrieved from *ensemble.lepbase.org* (http://ensembl.lepbase.org/index.html) as of Sept 1 2019, with the exception of the search for almost-universal orthologs in different assemblies, which was done here with the same sets of query genes.
† Using complete gene evidence only. CEGMA: C, complete; P: partial. BUSCO: C, complete (uni and multicopy); D, duplicated or multicopy; F: fragmented. The number of complete unicopy BUSCOs can be calculated as the difference between the total number of complete BUSCOs and the number of multicopy BUSCOs.

### Comparison to other *D. plexippus* genome assemblies

Although the assemblies DpMex_v1, Dpv3, and its derivative Dpv4 (7,11), have virtually the same span (∼249 Mb), all of them are still smaller than the genome size of 284 Mb estimated by flow cytometry (84). A plausible explanation is that all three assemblies reflect reasonably well the euchromatic but not the heterochromatic genome portion, as the latter requires specialized approaches for sequencing due to its repetitive content (86). Particularly heterochromosome *W*, one of the two exceptionally large chromosomes of *D. plexippus* along with heterochromosome *Z*, is misrepresented in the current and previous assemblies as at least two thirds of its length are highly heterochromatic (9).

The DpMex_v1 assembly shows a highly improved contiguity compared to Dpv3 (contig number: 108 vs 10,682) and the N50 increases from ∼111 kb to ∼3,941 kb, which results essentially from the increased length of the PacBio sequencing reads used. Relative to the Dpv4 assembly (11), which implements Hi-C to scaffold Dpv3, the scaffold N50 metric is 1 Mb lower (8.16 vs 9.21 Mb) but heterochromosome *Z*, which is represented by the largest scaffold, is 727 kb longer in DpMex_v1 (16.34 Mb vs 15.62 Mb; chromosome *1*, see below) (Fig. 1A). Unlike Dpv4, DpMex_v1 includes only 23 chromosome-length scaffolds (see below). Overall, the high contiguity of both DpMex_v1 and Dpv4 places both assemblies together with that of *H. melpomene* among the few lepidopteran genomes with multi-megabase N50 (Table 1).

**Figure 1.**
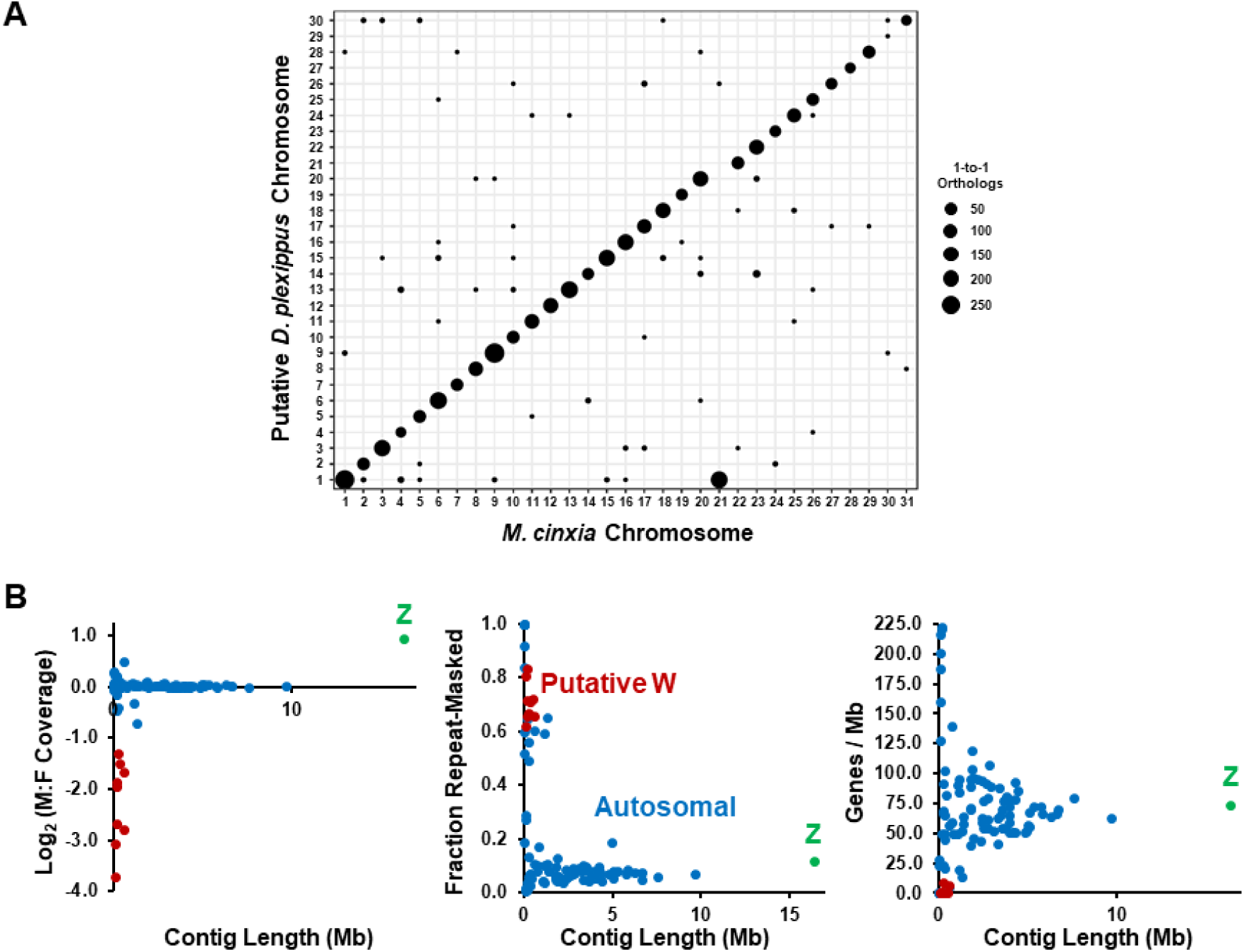
Assignation of the DpMex_v1 genome assembly to the chromosomes of *D. plexippus*. (A) Chromosomal anchoring of contigs based on the chromosome of *M. cinxia* showing the highest number of one-to-one orthologs for protein-coding genes in our OGS1_DpMex. 97.02% (4,855 out of 5,004) orthologs mapped to chromosomes of *M. cinxia* provided coherent cytological information about to which chromosomes the contigs should be anchored. The diameter of the circles denotes the number of such one-to-one orthologs. Overall, high synteny conservation between the two species can be observed which supports that the gene content of the ancestral chromosomal elements to these two species remained essentially intact. It can also be observed that chromosome 1 in *D. plexippus* encompasses genes from chromosomes 1 and 21 of *M. cinxia*, reflecting the outcome of a fusion event. (B) For each of the 108 contigs part of the DpMex_v1 assembly, the log_2_ of the male to female coverage (left), the repeat-masked fraction (center), and gene density (right), is plotted against its length. A color code is used to distinguish the contigs categorized as Z-, autosomal, - and potentially W-related.

As for the agreement in internal order and orientation of scaffolds, we detected 131 differential sequence discontinuities in Dpv3 relative to DpMex_v1 affecting 61 Dpv3 scaffolds (Supplemental Table S3). Importantly, 3 of these discontinuities were already reported as missassemblies of Dpv3 (9). In addition, DpMex_v1 has a similar span as Dpv3 and Dpv4, but virtually no gaps while 2.3% of Dpv3 and Dpv4 correspond to gaps. Further, relative to Dpv4, the two genomes exhibit a high degree of upper-level collinearity so that large-scale misassemblies in any of them is unlikely (Fig. S7). Nevertheless, at a finer scale, multiple small differences were uncovered. At this time, it is not apparent what is the relative contribution of *bona fide* genetic differences between the individuals sequenced and misassembly errors. The differences observed in the K-mer spectra between both assemblies (Fig. S8), particularly the underrepresentation of high multiplicity K-mers, *i*.*e*. those less likely to represent sequencing errors, which are present in the sequencing data generated for the DpMex_v1 assembly and absent in Dpv3, is striking.

Lastly, DpMex_v1 exhibits increased gene completeness compared to Dpv3 and Dpv4, and substantially higher than that of for example *H. melpomene* v2.5, according to the BUSCO analysis. The completeness was similarly high according to CEGMA, which uses a different reference gene set (Table 1). Specifically, the DpMex_v1 assembly harbors more unicopy complete (54 and 55) and fewer (51 and 58) duplicate BUSCOs than Dpv3 and Dpv4, respectively. Through TBLASTN, we confirmed the presence of significant hits for 48 of the 57 missing BUSCOs, including the 20 BUSCOs undetected across all three *D. plexippus* assemblies (Supplemental Table S4). These were also missed in the genome assembly of *H. melpomene* v2.5, confirming that the DpMex_v1 assembly is the most complete among those available in *D. plexippus* (Supplemental Text S3 and Tables S4-S5).

### Repeat and gene annotation

To delineate transposable element (TE) insertions and low-complexity repeat sequences, we first performed an *ab initio* repeat modelling step, which was followed by masking the final, polished genome assembly, including low-complexity regions and simple repeats. In total, ∼42.9 Mb (17.26%) were masked. A categorization of the mentioned repeats shows that 19.25 Mb (7.75%) correspond to interspersed repeats (Supplemental Table S6).

A new gene annotation (OGS1_DpMex) was generated for the DpMex_v1 assembly by considering different types of support: (*i*) RNA-seq data from 14 different types of biological samples from larval, pupal, and adult stages (see below); (*ii*) by identifying a homolog in at least one of six other lepidopteran species or *D. melanogaster*; and (*iii*) by having an equivalent gene model in the previous annotation OGS2 (7). The new annotation includes models for 15,995 protein-coding genes (Table 2; Fig. S9A), with 7.1% of the gene models being supported only by RNA-seq data, 7.4% only by homologous sequences, and 82.23% supported by both. Three quarters of the remaining 3.3% of the gene models are supported by having an equivalent, although not necessarily identical, counterpart in OGS2 (Fig. S9), with the rest being purely *de novo* computational predictions.

**Table 2.**
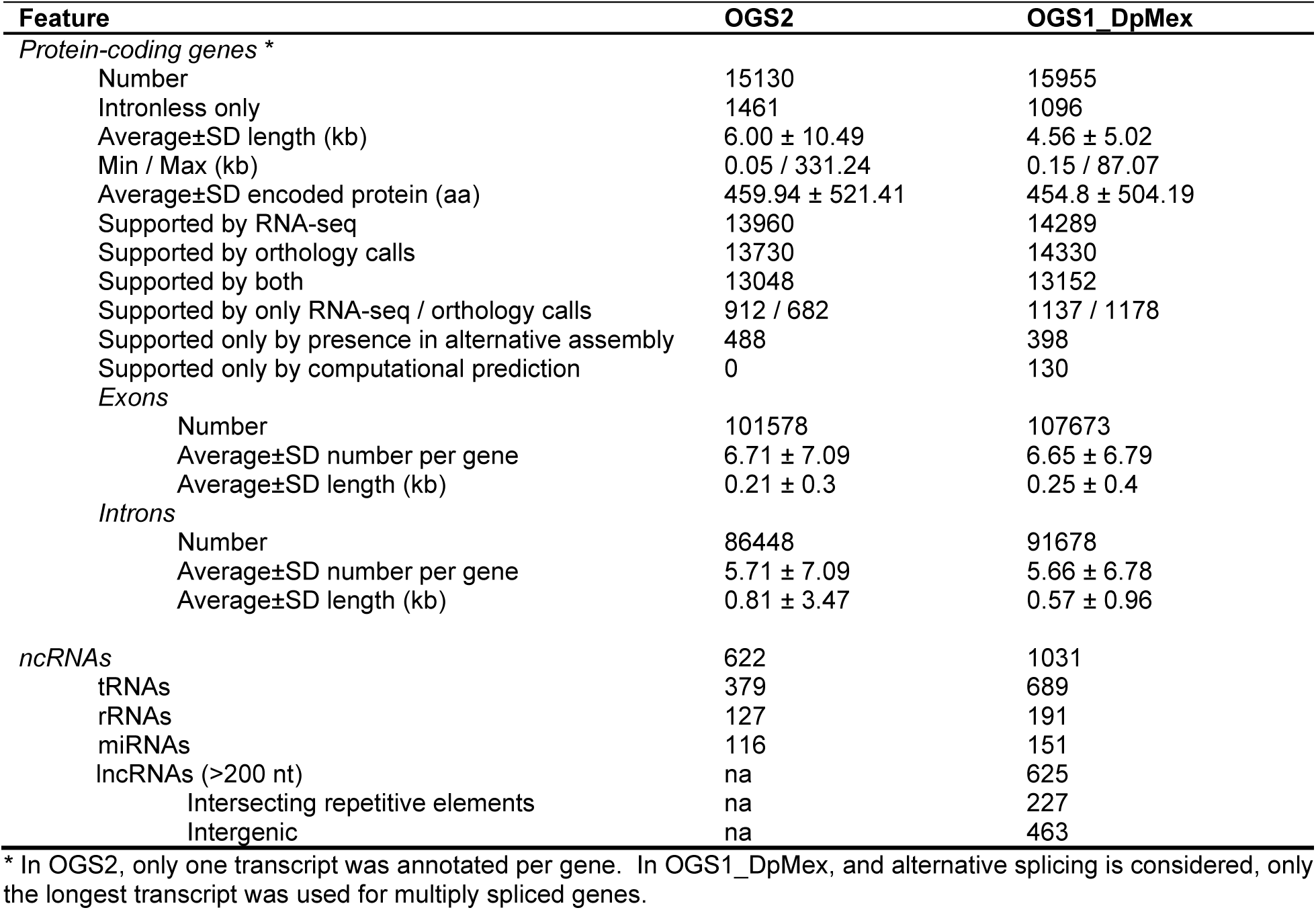
Salient features in a previous and new gene annotation of *D. plexippus*.

Sequence similarity searches using proteomic data from six other Lepidoptera revealed support by homology in at least one of the species considered for 14,330 (89.6%) of the gene models (Table 2 and Supplemental Table S7). The direct comparison between the annotations of protein-coding genes revealed that 1,077 (7.1%) of OGS2 were not found in OGS1_DpMex while 1,677 (10.5%) gene models of OGS1_DpMex showed the opposite trend. Among these sets of gene models unique to OGS2 and OGS1_DpMex respectively, the proportion of those with homology support is also higher for the latter (437 or 40.6% vs 834 or 49.7%). Importantly, when we compared the absolute number of gene models from both assemblies for which it was possible to find at least one homolog with OrthoFinder (87), we found a higher number of matches for the OGS1_DpMex gene models regardless of the species (Fig. S10), reaching a maximum difference between OGS1_DpMex and OGS2 in the case of *H. melpomene* with 585 additional gene models in OGS1_DpMex.

Among the gene models found in both OGS2 and OGS1_DpMex, 8,861 correspond to single copy gene entities while the remaining 2,411 gene entities appear in the form of two or more copies in at least one of the annotations. Among the latter, 664 gene entities show exactly the same number of copies in OGS2 and OGS1_DpMex (1556) while the other 1,747 gene entities differ in copy number between assemblies, with a net difference of 313 (3901 – 3588) for OGS1_DpMex. Overall, OGS1_DpMex shows an increase in the number of multi-copy gene entities, including those that are single copy in OGS2 (2-sample test for equality of proportions, Χ^2^= 6.164, d.f. = 1, *P =* 0.013). These differences can represent an overall more accurate assembly, true differences in copy number between the individuals sequenced, or fragmented predicted models. An extreme example corresponds to a gene entity that exhibits significant sequence similarity by BLASTP against the protein-coding gene Cation-Channel complex subunit UNC-79 from the butterfly *Vanessa tameamea*. This gene entity is single copy in OGS2 whereas it is present in 25 copies in OGS1_DpMex scattered through 16 contigs.

Regarding ncRNA genes, our new annotation includes 1,656 gene models, *i*.*e*. a 2.7-fold increase relative to OGS2 (Table 2). The increase is consistent across all categories of small ncRNA genes, *i*.*e*. tRNAs, rRNAs, and miRNAs. Additionally, we used ribodepleted stranded RNA-Seq of pooled samples spanning the larval, pupal, and adult stages to help predict lncRNA genes, which were omitted in OGS2, via a dedicated pipeline. These genes were required to generate transcripts longer than 200 nt, encompass at least one splicing junction, and be antisense if overlapping with a protein-coding gene model. Under conservative criteria, we annotated 625 lncRNA gene models (Table 2), 463 of them being intergenic, 134 being antisense to coding sequences, and 28 residing in introns of models of protein-coding genes. As in humans and other insects, a sizable number of the lncRNA models (227 or 36.3%) overlap with TE sequences (88,89).

In summary, the new gene annotation OGS1_DpMex encompasses 17,651 models, *i*.*e*. 1,899 more than OGS2, of which 865 correspond to protein-coding genes and 1034 to ncRNA genes (Fig. 9B). Collectively, one third of the genome (32.1% or 78.5 Mb) is transcribed into primary transcripts with 10.8% being associated with mature transcripts, and CDS sequences representing 8.9% of the genome.

### Genome assembly assignment to chromosomes

Contig anchoring to the *D. plexippus* chromosomes was performed assuming a high level chromosome conservation of gene content, *i*.*e*. macrosynteny, which is heavily supported by comparative analysis involving *M. cinxia, H. melpomene*, and other Lepidoptera (9,90-93). [*M. cinxia, H. melpomene*, along with *D. plexippus*, belong to the Nymphalidae family, the largest among the butterflies (90,94)]. Accordingly, each DpMex_v1 contig was anchored to the *M. cinxia* chromosome that harbored the highest number of 1-to-1 orthologs between protein-coding genes of the OGS1_DpMex and those of *M. cinxia*. This species presumably preserves the ancestral lepidopteran karyotype n = 31 (94,95), being phylogenetically close to *D. plexippus* (9). The anchoring process was based on positional information from 5,004 1-to-1 orthologs, which resulted in 74 out of 108 contigs being anchored directly to chromosomes (Fig. 1A; Fig. S12A; Supplemental Table S8), and 6 more indirectly as they are scaffolded with some of the former using RaGOO. Importantly, these contigs span ∼238.4 Mb, *i*.*e*. 97.2% of the total assembly length, and include 97.02% (*i*.*e*. 4,855) of the 1-to-1 orthologs mapped. The remaining 149 1-to-1 orthologs could have been involved in interchromosomal gene transposition events.

All but chromosomes *3, 9, 13, 16, 19, 21*, and *28* are represented by a single scaffold. Crucially, all contigs of the same scaffold, if mapped, agreed in their chromosomal assignment, highlighting the reliability of the strategy followed. The largest chromosome spans 16.34 Mb and includes genes from chromosome *1* and *21* of *M. cinxia*, confirming a previously inferred fusion event that predated the radiation of the genus *Danaus* (9)(Fig. 1A). Such chromosomes correspond to the ancestral- and the neo-portion of the heterochromosome *Z* of *D. plexippus*, respectively (9,94). Like chromosome *1*, chromosomes *15, 22, 23*, and *30* are also associated with a single contig and scaffold.

To substantiate further our chromosome anchoring process, we used two approaches. First, we reasoned that if the overall macrosynteny conservation assumption was correct, the majority of the gene content in the chromosomes of *D. plexippus* chromosomes should also be found in species such as *H. melpomene* and the silkmoth *B. mori* –a representative of the moths and whose ancestor split from that of butterflies during the Cretaceous period (96). Based on 1-to-1 orthologs between the indicated species and *D. plexippus*, we corroborated the presumed microsynteny conservation (Supplemental Text S4 and Fig. S14). Second, to confirm the distinction between homo- and heterochromosomes, we calculated the average log_2_ average male to female coverage for every contig by mapping Illumina genomic DNA sequencing data from 70 libraries corresponding to 36 males and 34 females (6)(Fig. 1B and Supplemental Fig. S13 and Table S9). A twofold difference in coverage, *i*.*e*. a log_2_(M:F)=1, is expected for the unicopy sexual chromosome in the heterogametic sex, *i*.*e*. the *Z* chromosome in females (*ZW*), but not for the autosomes (9,97). The contig corresponding to chromosome *Z* stands out as it is the only contig with a log_2_ ratio close to 1. Nine additional contigs (Supplemental Fig. S12B), none of them anchored to any chromosome, exhibit log2 values < -1, which opens the possibility that they can be components of the *W* chromosome. These contigs are mainly composed of genome-wide repeats, are relatively short (<0.59 Mb; median size 0.14 Mb), and exhibit very low gene densities; in fact, 7 of the contigs harbor no annotated gene (Fig. 1B; Supplemental Table S10). With the exception of chromosome *W*, these results strongly support the reliability of the anchoring process of our genome assembly to the chromosomes of *D. plexippus*.

### Transcriptome Atlas

We sequenced over 926 million strand-specific paired-end (PE) reads resulting in 157 Gb of sequence data. Most of these RNA data corresponds to poly(A)+ transcripts from 14 different types of biological samples, including larval stages, pupal stages, and anatomical parts of 2-day-old posteclosion individuals (Fig. 2A). For all biological sample types, we obtained biological replicates from two different individuals, except for 1^st^ instar larvae and adult heads, for which replicates were made from pools to obtain sufficient RNA (see Material and Methods). Additionally, we incorporated our non-poly(A)+ enriched RNA-seq data from the four types of pooled samples. In total, our transcriptome atlas is based on 36 distinct samples. On average, 87.4% of the reads from each sample mapped to our genome assembly, with 49.2% mapping to unique locations and 38.2% mapping to multiple locations. Ultimately, 76.7% of all sequencing reads could be confidently assigned to the annotated fraction of the assembly. Reads that only mapped to introns (2.4%), rRNA (32.7%) and tRNA (<0.1%) were omitted in downstream analyses, keeping only the 41.6% of reads that mapped to protein-coding, lncRNA, and miRNA genes (Supplemental Table S11).

**Figure 2.**
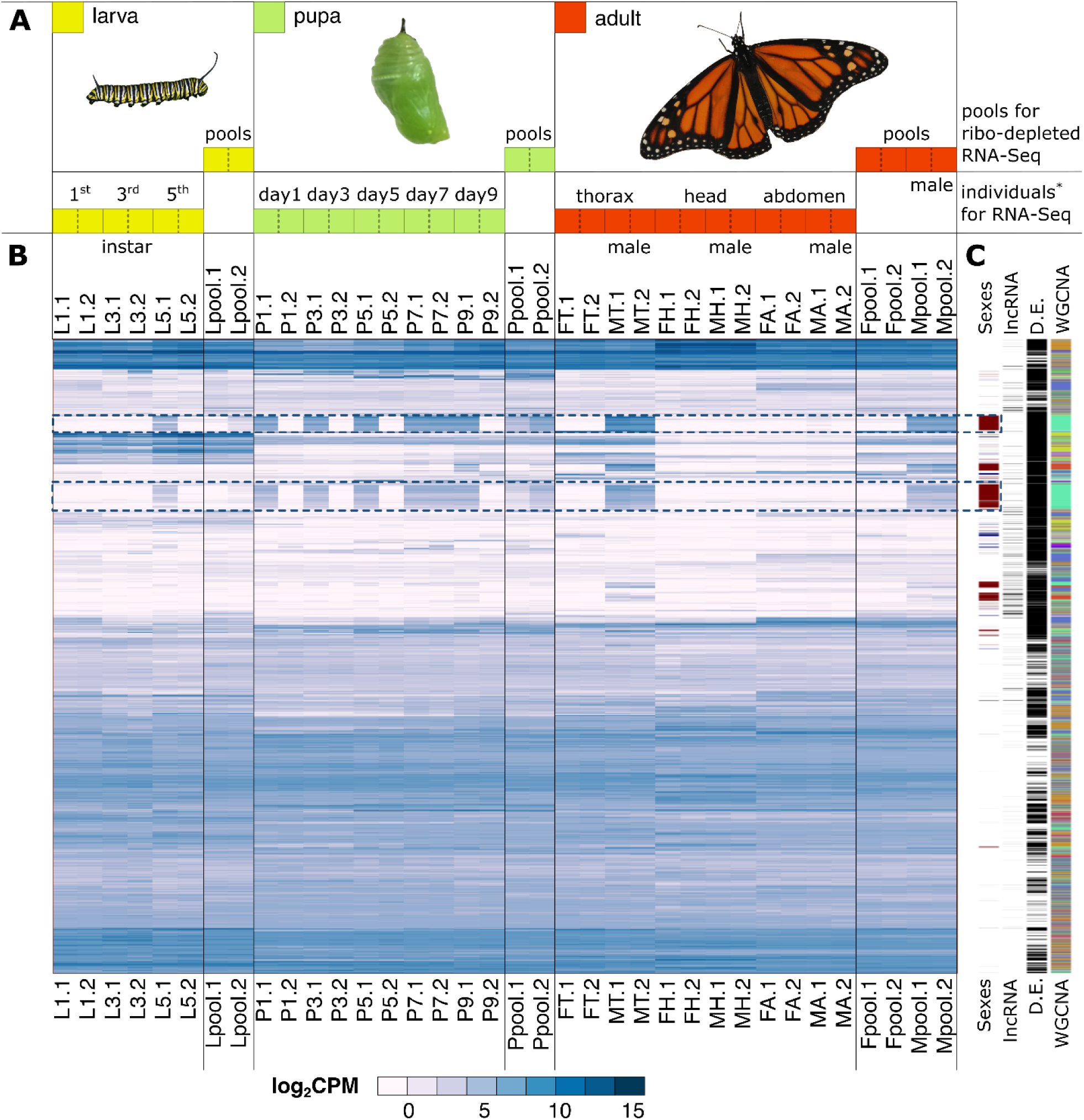
Transcriptome atlas experimental design. (A) Fourteen specific stages and anatomical parts were RNA-sequenced. These include: 1^st^, 3^rd^ and 5^th^ instar larvae (yellow boxes); day 1, 3, 5, 7 and 9 for pupae (green boxes); and thorax, head, and abdomen from 2-day-old posteclosion males and females (red boxes). Two biological replicates of these 14 sample types were sequenced to an average of 20.7 million PE reads each. In addition, total RNA from four additional sample types corresponding to broadly defined developmental stages (larva, pupa, adult males and females) were ribodepleted and sequenced to an average of 43.3 million PE reads. These samples consisted of pools of individuals of the same stage, for which two biological replicates were also included. The two replicates for each of the 18 sample types are indicated by the .1 or .2 at the end of their name. (B) Heatmap of library-size normalized gene-level log_2_CPM of 14,865 expressed genes (rows) across 36 samples sequenced (columns). Dotted boxes highlight two groups of genes with marked male-biased expression. (C) From more internal to more external, selected gene categories: genes with sex-biased expression (male-biased dark red, female-biased dark blue); lncRNA genes; D.E., differentially expressed genes in at least one contrast; WGCNA, genes clustered according to this methodology.

After gene-level quantification and normalization (see Material and Methods), we deemed genes as expressed when showing ≥1 count-per-million (CPM) in a sample. We found 7,479 genes expressed in all 36 samples, with this number progressively increasing as we required expression from fewer samples, reaching a total of 14,865 genes detected in at least two samples (Supplemental Fig. S14 and Table S12). Across different types of biological samples, we detected substantial differences in the number of expressed genes, with the four head samples featuring the lowest (9,013 - 9,822) and the two whole-body male pooled samples featuring the highest (13,180 and 13,309) number of expressed genes (Supplemental Fig. S15). Further, Multidimensional Scaling (MDS) analysis largely corroborated the developmental relationship among the sequenced samples and the replicate consistency (Supplemental Fig. S16A), while revealing the marked signature of sex-biased expression on the global profile, which is also reflected as a distinctive trend shown by particular sets of genes (Fig. 2B-C, dotted boxes). Interestingly, these gene sets exhibit the same sex-specific bias in thorax but not in head or abdomen, which could plausibly result from a comparatively delayed sexual maturation of those anatomical parts in relation to thorax. Once the sex-biased genes were omitted, MDS grouped all samples consistently according to their stage, and biological replicates were also positioned systematically closer (Supplemental Fig. S16B). Although larval samples did not perfectly separate by instar, the pupa samples followed a consistent bidimensional trajectory, starting at day 1 (P1) and ending at day 9 (P9).

### Expression patterns across the transcriptome atlas

We next sought to investigate differential gene expression across developmental stages and adult anatomical parts. We avoided performing all possible pairwise comparisons involving the 18 types of biological samples profiled (*i*.*e*. 153 comparisons) and, with a few exceptions, we primarily focused on which genes in a given sample type were differentially expressed in relation to other types of samples from the same stage (Supplemental Tables S13 and S14). To deem a gene as differentially expressed, we required a significantly higher than 2-fold expression difference at a 5% false discovery rate (FDR). Upon omitting those genes that only showed differences in mRNA abundance in our technical contrast (*Source* in Supplemental Table S13), 9,497 genes –63.9% of all genes expressed– showed statistically significant differential expression in at least one of the contrasts performed, with 1,561 (10.5%) only in one. The contrast corresponding to the transition between larva and pupa (P1:L5) displayed the largest number of expression changes (Supplemental Table S13 and Fig. S17).

As *D. plexippus* oviposition occurs in milkweed host plants that contain toxic cardenolides, larval instars are crucial in the context of host adaptation, particularly because sequestration of cardenolides is higher during early than late instars (98). Therefore, transcriptome characterization of larval instars is vital to understand how gene function changes in the context of for example invasive species (99). In total, up to 2754 genes are differentially expressed in at least one of the contrasts involving larval instars although the number of such genes changed substantially (Supplemental Fig. S17 and S18). For example, in the contrast *L1:L*, which entails the comparison of L1 to the average of L3 and L5 (Fig. 3), we identified 867 genes as upregulated and 574 as downregulated in L1 (Fig. 3A-B and Supplemental Table S13). In the remaining contrasts within the larval stage, such numbers are 596 and 600 (*L3:L*), and 1019 and 421 (*L5:L*), respectively (Supplemental Fig. S18). Although there is some overlap in the identity of the differentially expressed genes in these three contrasts, many of them are different. This is apparent when examining the patterns of the differentially expressed genes in one of the contrasts in the context of the remaining samples (Fig. 3C-D; Supplemental Fig. S18). For instance, L1 upregulated genes are also highly expressed in abdomen samples of both sexes, while L3 and L5 upregulated genes are not. In this case, although we cannot discard some influence by early sexually-differentiated genes, the observed pattern could represent a second wave of differential expression during adult tissue differentiation as observed in *D. melanogaster* (100). Functional enrichment analyses also confirmed the differential overrepresentation of GO terms of the Biological Process ontology (*P*_adj_ < 0.2) across the three contrasts. For example, genes significantly upregulated in L1 relative to L3 and L5 appear to be preferentially related to signal transduction, neurotransmitter transport, and cell communication, while those downregulated were largely to metabolic processes and immune response. These trends are compatible with a relative metabolic activation once the caterpillars start to feed and grow (thus lower expression in L1 but higher in L3 and L5), which is concomitant to a reduction of signaling processes required for very early development (Supplemental Fig. S18 and Table S15). Importantly, when we compared all larval instars to the remaining individual samples (pupa and adult parts), we detected a significant enrichment for members of gene families implicated in detoxification. Thus, we found 74 genes encoding proteins that include a Major Facilitator Superfamily domain and 34 containing a Cytochrome P450 domain, while 14 genes encoded UDP-glucosyl transferases. When we analyzed the individual larval instars, Cytochrome P450 genes are relatively depleted in L1 but enriched in L5, highlighting further the differential transcriptome deployment of gene functions across larva instars.

**Figure 3.**
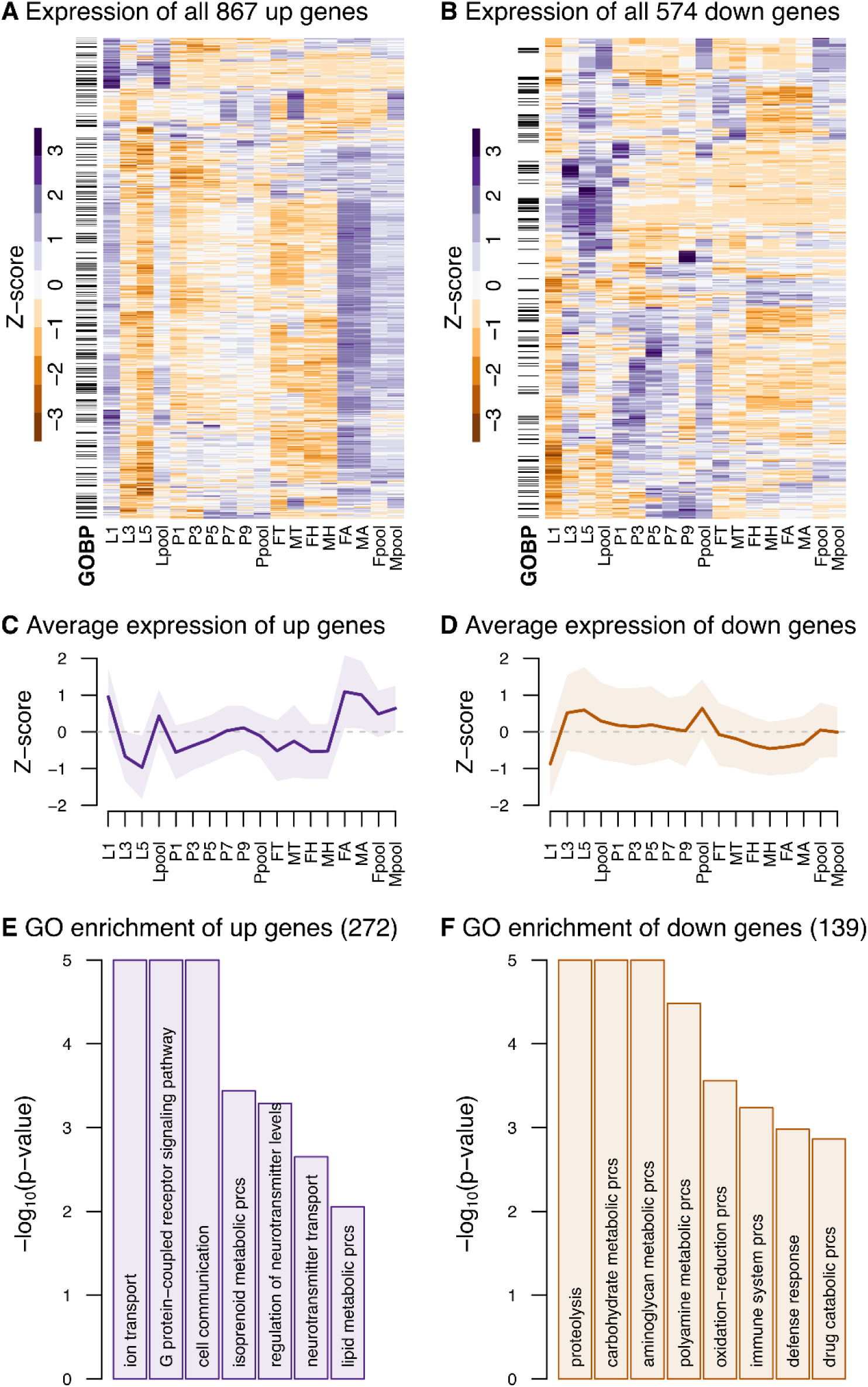
Differential expression results for the L1:L contrast. (A, B) Heatmaps of differentially upregulated and downregulated genes (5% FDR) in L1 relative to the average of L3 and L5. The average of each pair of replicates is used for the columns, and the rows are scaled using a Z-score. (C, D) Average gene expression trend based on the Z-score scaled expression of all upregulated and downregulated genes across all samples, respectively. The shaded area represents ±1 SD from the mean. (E, F) Topmost significantly enriched Biological Process GO terms amongst upregulated and downregulated genes, respectively.

With the end of the larval stage, the development of adult body structures is the primary process during pupation, which can be impacted by the effects of parasites such as *Ophryocystis elektroscirrha* (101). We predicted that expression patterns would differ markedly from those in larvae, as well as between early and late pupae. In good agreement, the contrast *P1:L5* revealed the upregulation in P1 of genes related to developmental and signaling pathways. Likewise, the contrasts *P1:P* and *P9:P* differ substantially in the functional attributes of the upregulated and downregulated genes. For example, while immune system and defense response are strongly associated to upregulated genes in *P1*, these appear not only to be downregulated in *P9* but replaced by others related to the nervous system and a wide variety of metabolic processes (Supplemental Fig. S18 and Table S15). In total, a minimum of 969 (*P3*:*P*) and a maximum of 2628 (*P9*:*P*) genes were found to be differentially expressed across the contrasts involving pupa samples (Supplemental Table S13), while 3943 genes were differentially expressed in at least one of the contrasts involving pupa related sample types (Supplemental Fig. S17).

Lastly, and to shed some light into the 5,090 genes that were not called differentially expressed in any contrast, we applied Weighted Gene Correlation Network Analysis (WGCNA) to all genes expressed (102). Twenty-five clusters were delineated (Supplemental Table S14 and S16), with some of them corresponding quite tightly to particular sets of differentially expressed genes (*e*.*g. clust16* and *Sexes*.*up*; Fig. 2C), while others overlapped with different sets of differentially expressed genes (*e*.*g. clust5* and *clust9*). Notably, this complementary clustering approach also uncovered expression patterns that were overlooked in the differential expression analysis. For instance, *clust06* included 684 genes that progressively diminish their expression during all larva and pupa stages and are further diminished in the adult samples (Supplemental Fig. S19). These genes are enriched in GO terms related to ncRNA metabolism, RNA processing and DNA replication, and PFAM RNA-binding domains. Another cluster, *clust08*, consists of 527 genes that follow almost the opposite pattern: increasing during development and reaching their highest level in the adult samples (Supplemental Fig. S19 and Supplemental Table S15). Interestingly, these genes show few signatures of common functionality, except for those involved in ion transport and the structure of insect cuticle. This cluster contains a relatively small fraction of genes with a GO annotation (150/527 for Biological Process), providing an opportunity to study genes with new functions. Together, the patterns reported represent the most comprehensive dynamic portrait of gene expression so far generated during the life cycle of *D. plexippus*.

### Sex-biased gene expression

We found 14.42% (2,143/14,865) differentially expressed genes between the sexes in at least one of the four types of adult samples assayed (Supplemental Table S13; Fig S20), which is roughly half of the one third documented in *D. melanogaster* when a 2-fold cutoff is applied (103). This could be in part the result of differences in statistical power and criteria used. We also observed very limited overlap among sample types, denoting differences in tissue composition and potential to harbor sex-biased expression. In species with heteromorphic sexual chromosomes, sex-biased expression can result from different factors including gene presence in only one sex (*i*.*e*. when residing on the W, in female Lepidoptera) and from gene presence on the heterochromosome that is diploid in the heterogametic sex (*i*.*e*. on the Z, in male Lepidoptera) coupled with incomplete dosage compensation, *i*.*e*. absence of whole-chromosome upregulation of the heterochromosome present in one copy in the heterogametic sex (*i*.*e*. the Z in female Lepidoptera) (104,105). Recently, a comparison of the brain transcriptome between adult males and females documented a difference in dosage compensation between the anc*-Z* and neo*-Z*. The anc*-Z* showed roughly half of the expression of the autosomes in both males and females due to downregulation in males while the neo*-Z* showed almost equal level of expression relative to the autosomes in both sexes through a newly evolved upregulation in females (11). Here, we examined the reproducibility of such patterns across the adult sample types we assayed.

First, we compared the median absolute expression Z:A ratio within each sex finding both commonalities and differences across sample types relative to the reported pattern (11) (Fig. 4). For female whole-bodies and individual anatomical parts (heads, thorax, and abdomen), we observed a significantly lower median expression level for the anc*-Z*, but not for the neo*-Z*, relative to the autosomes, corroborating first the lack of complete dosage compensation for the anc*-Z*, and second the newly evolved complete dosage compensation for the neo*-Z* in this sex. Nevertheless, the median expression ratio of the anc*-Z* compared to the autosomes goes from nearly half in heads to closer values to 1 (maximum = 0.82, whole-bodies), denoting that absolute lack of dosage compensation happens only in heads. In contrast, for males, the presumed repression of the two Z chromosomes that should result in also a significantly lower expression level relative to the autosomes is observed in heads and, to some degree, in abdomen as reported in other Lepidoptera (106-108) but not for thorax and whole-bodies, suggesting that this pattern is likely tissue-dependent and therefore obscured in those sample types in which the repression mechanism does not predominate. Importantly, these patterns are robust across several expression thresholds (Supplemental Fig. S21) and are not the result of collapsing all the autosomes (Supplemental Fig. S22).

**Figure 4.**
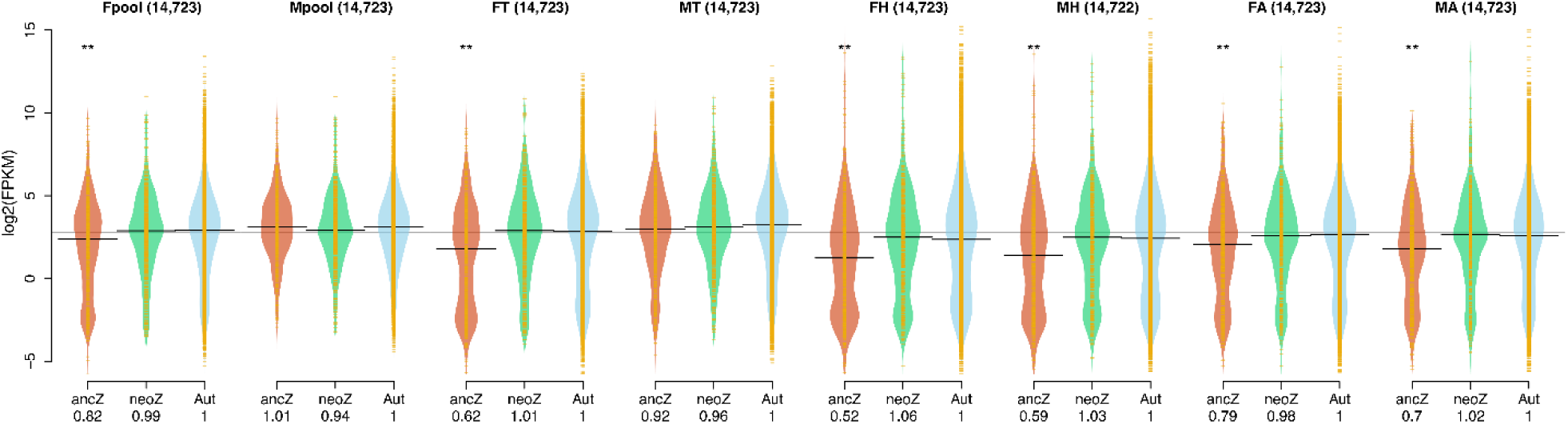
Whole-chromosome expression in females (ZW) and males (ZZ) in four sample types of *D. plexippus*. Sampled assayed: pools of whole-body males and females (Mpool, Fpool); individual samples of thorax (MT, FT), abdomen (MA, FA), and heads (MH, FH). The bean plots show the distribution of absolute normalized log_2_ expression values in FPKM for the ancestral (anc-) and neo (neo-) portions of chromosome 1 (i.e. the Z chromosome) and the autosomes. The horizontal line in each plot corresponds to the median expression value. A global median value across sample types is shown with a grey line in the background. The median Z:A ratios are shown at the bottom. For each sample type, statistical significance was established according to Wilcoxon signed-rank tests and upon applying a Bonferroni correction (** *P*<0.01). The number of genes considered is indicated on top of each bean plot. A minimum expression threshold of 0.01 FPKM was required. The results with other thresholds and for the autosomes considered separately can be found in Supplemental Fig. S21 and Fig. S22 respectively.

Next, we examined the median expression ratios between females and males for the autosomes (AA:AA) and the two portions of the *Z* heterochromosome (anc*-Z*:anc*-Z*anc*-Z*; neo*-Z*:neo*-Z*neo*-Z*) to assess the degree of expression equalization between the sexes, which is determined by the degree of dosage compensation of the *Z* heterochromosome in females and the downregulation of the *Z* heterochromosome in males (Supplemental Fig. S23 and S24). In good agreement with the observations above, the female to male ratios for the anc*-Z* are no different or slightly -but significantly-lower to those for the neo*-Z* and the autosomes in abdomen and head. In contrast, for the whole-body and thorax samples, the female to male ratio for the anc*-Z* shows only partial evidence of equalization between the sexes, being significantly lower. Only in whole-body samples, the neo*-Z* shows a slightly -but significantly-lower female to male ratio relative to the autosomes. Collectively, all these results underscore their sample-dependent nature while highlighting that dosage compensation is either absent or incomplete in the ancestral portion of *D. plexippus Z* chromosome, which is also reflected on the lack of perfect expression equalization between the sexes, this last pattern less acutely detected for the neo portion of the *Z* chromosome.

Lastly, we examined the chromosomal distribution of sex-biased genes across the autosomes, the ancestral*-Z*, and the neo*-Z* in whole-body and thorax samples as those are the ones with more sex-biased genes (see above) and therefore we have more statistical power (Supplemental Table S17). Based on the incomplete lack of dosage compensation featured by the anc*-Z*, we predicted an enrichment of male-biased genes for this portion of the *Z* heterochromosome but not for the neo*-Z*, in good agreement with previous observations in other Lepidoptera (108-110). Nevertheless, some interspecific variation relative to this pattern of enrichment has been observed (107), underscoring the influence of other factors, mainly sexually antagonistic selection (111,112), which might or might not align with the expectation based on the lack of complete dosage compensation (111,112). For whole-body, we found statistically significant evidence of a global non-random distribution of male-and female-biased genes across the different portions of the heterochromosome *Z* and the autosomes, a pattern robust across different thresholds of minimum expression (Cochran-Mantel-Haenszel test; whole-body, Χ_MH_^2^=56.20, d.f.=2, *P=*6.27×10^−13^; Supplemental Table S17). Analysis of the adjusted standardized residuals (113) confirmed that the anc*-Z* exhibited enrichment for male-biased genes and depletion for female-biased genes, with the autosomes harboring comparatively a significantly lower proportion of the first gene category and a higher of the second. The neo*-Z* showed no bias of any kind. For thorax, support for the same global non-random distribution is found (Cochran-Mantel-Haenszel test; thorax, Χ_MH_^2^=21.87, d.f.=2, *P=*1.79×10^−5^), although its statistical significance did not hold when the different thresholds of minimum expression were examined, arguably due to a more limited statistical power (Supplemental Table S17). Overall, our results adhere to the expected enrichment for male-biased genes in expression in the portion of the heterochromosome *Z* (anc-*Z*) that shows incomplete dosage compensation (107,108). The neo-*Z* portion not only does not show the same pattern but also exhibits a significantly lower global fraction of sex-biased genes, in fact, even lower than the autosomes (Cochran-Mantel-Haenszel test; whole-body, anc-*Z* vs neo-*Z* vs A: Χ_MH_^2^=185.08, d.f.=2, *P=*2.2×10^−16^; neo-*Z* vs A: Χ_MH_^2^=20.84, d.f.=1, *P=*5.00×10^−5^), which holds across thresholds of minimum expression (*P*_adj_*<*0.05 for each threshold; Supplemental Table S18).

### Long non-coding RNAs

Owing to the limited functional characterization of lncRNAs beyond model organisms (114,115) and the limited possibility to transfer information from other species due to their poor sequence conservation (116), we explored the expression patterns of lncRNAs throughout our transcriptome atlas. As in other species (115,117,118), we found that lncRNAs are expressed at a significantly lower level during the life cycle of *D. plexippus* (Supplemental Table S19), and exhibit more restricted expression profiles than protein-coding genes according to the Yanai index (Wilcoxon rank-sum test, *P=*2.2×10^−16^ ; Supplemental Fig. S25). The 492 lncRNA gene models (78.7% of the 625 annotated) found expressed with at least 1 CPM in at least two samples of our atlas did not fall randomly across our sets of differentially expressed genes or WGCNA clusters (Fig. 2C). Intriguingly, three clusters (*clust10, clust22* and *clust23*) contained a higher number of lncRNAs than expected (one-tailed Fisher’s exact test, *P*_adj_ < 0.05; Supplemental Table S16). Likewise, certain sets of differentially expressed genes were also prone to harbor more lncRNAs than expected (Supplemental Table S13). For instance, the contrast *Source*, which compares pools of larva, pupa and adults of both sexes to those from individual developmental stages and adult anatomical parts, yielded the highest enrichment. This was expected as the library preparation for transcriptome sequencing of the pooled samples purposely enriched for lncRNAs by minimizing loss of non-polyadenylated transcripts.

Interestingly, the contrast showing the second largest number of differentially expressed lncRNAs was the comparison of pooled whole-body adult females compared to males (contrast *Sexes*, Supplemental Table S13). Among the differentially expressed genes in such contrast, and relative to protein-coding genes, we did not detect differences in the proportion of lncRNAs genes between females and males (20/245 vs 81/1563, two-tailed Fisher’s exact test, *P=*0.071). In sharp contrast, however, and again relative to protein-coding genes, we did detect a significant increase of differentially expressed lncRNAs when we compared pooled male and female adults against pooled larva and pupa (contrast *Adulthood*, Supplemental Table S13; 71/1244 vs 33/1277, two-tailed Fisher’s exact test, *P=*8.2×10^−5^). Thus, while 21.13% (101/478) of the lncRNAs shows statistically significant sex-biased expression, only 11.99% (1707/14233) of the protein-coding genes do (2-sample test for equality of proportions with continuity correction, Χ^2^=34.97, d.f.=1, *P=*3.35×10^−9^).

Further examination of the GO terms associated with the protein-coding genes enriched in the same clusters as lncRNA genes allowed the tentative functional categorization of the latter. For example, *clust23* includes a low number of genes but harbors the highest fraction of lncRNAs (21/58, 36%), showing high expression in most of the pooled samples. Interestingly, *clust23* shows increased expression during larva development, and higher expression in heads compared to thorax. Although only 5 genes have an annotated GO Biological process, there is an enrichment for nitrogen compound transport and exocytosis terms (q-value of 0.03 and 0.04, respectively). In summary, lncRNAs likely participate in essential biological process during the whole life cycle of *D. plexippus*, are more finely regulated during adulthood than during previous developmental stages, and are more heavily influenced by the differences between the sexes compared to protein-coding genes.

## CONCLUSIONS

The rapid transformation of the *D. plexippus* habitat has had profound consequences, including a population decline and a modification of migration patterns (3). This new ecological context for *D. plexippus* results in fewer or different oviposition opportunities (119), impaired adult and larval survival (120,121), and higher infection rates by parasites (122). A better understanding of the genetic basis underlying the adaptation process of *D. plexippus*, as well as effective conservation efforts, will require the use of genomic resources better representing the population genetic diversity of the species, as well as a more comprehensive knowledge about gene function and the regulatory basis of how the monarch transcriptome program changes during the life cycle.

The reference-quality genome assembly from a non-migrating colony resident at Mexico represents a step forward in this direction, mitigating the insufficiencies of depending on a single reference-quality genome assembly, including the presence of minor alleles for a set of loci, missing sequence, and the underrepresentation of genetic diversity at structurally dynamic regions (123-125). In addition, the new genome assembly and extended annotation generated here will prompt deeper comparative analyses of the gene organization across insects, revealing unique properties of the Lepidoptera chromosomes in general, and those of *D. plexippus* in particular, serving as a platform for future genome-wide population studies on structural variation, which is known to be particularly important in ecological adaptation across phyla (126-129).

Lastly, the portrait of the transcriptome program generated here serves as a baseline for the exploration of transcriptome responses associated to genotype-by-environmental interactions in the context of different host species (99,130-132). Monarch larvae have no choice of what they eat, spending multiple days on the plant chosen by their mothers for oviposition until they pupate. Understanding the developmental transcriptome provides a reference to understand the interplay between gene regulation and viability on alternative hosts, as well as taking into consideration lncRNAs, which are emerging as relevant adaptation drivers (16-18).

## DATA AVAILABILITY

Sequencing data were deposited to NCBI GenBank under the bioproject ID 663267, namely: Illumina DNA-seq data, SAMNX (tbd); Illumina RNA-seq data, SAMNY (tbd); SMRT sequencing data, SAMNT (tbd); and final genome assembly and annotation, NJBExxx.

## ACKNOWLEDGEMENTS

We thank Mahul Chakraborty for feedback on analytical procedures, and to Luis Herrera-Estrella and John Parsch for comments on early versions of this manuscript, respectively. We also thank the University of California, Irvine High Performance Computing cluster and Langebio’s Mazorka cluster, for facilitating most of the computational analyses, and the Secretaría de Medio Ambiente y Recursos Naturales (SEMARNAT), Subsecretaría de Gestión para la Protección Ambiental, Dirección General de Vida Silvestre, México, for granting us the sample collection permit OFICIO NÚM. SGPA/DGVS/02879/15.

## AUTHOR CONTRIBUTIONS

J.M.R, T.A.M, and C.A.G conceived and designed the experiments as well as wrote the manuscript. N.O.N and P.L.H were in charge of butterfly husbandry, sample collection, and total RNA extraction for subsequent sequencing. B.D.C extracted genomic DNA for subsequent sequencing. P.M.G generated the genome assembly and gene annotation. J.M.R and C.A.G performed most downstream analyses with further assistance in contig chromosome anchoring (P.M.G), WGCNA transcriptome network analysis (M.J.P.M, D.I.V, A.J.K), and lncRNA analysis (M.M.L).

